# Expression of Retroviruses in Guinea Pig Lymphomas

**DOI:** 10.64898/2026.01.13.699032

**Authors:** K. Brown, M. J. Dobromylskyj, H. A. Franks, A. Malecka, C. E. Staley, R. E. Tarlinton

**Affiliations:** Division of Virology, Department of Pathology, Addenbrooke’s Hospital, University of Cambridge, Cambridge, UK, CB2 1QP; The Veterinary Pathology Group, Unit 1b, Babbage Way, Science Park, Clyst Honiton, Exeter, UK, EX5 2FN; Histopathology Department, Finn Pathologists, One Eyed Lane, Weybread, Diss, Norfolk, UK, IP22 5TT; Centre for Cancer Sciences, Biodiscovery Institute, School of Medicine, University of Nottingham, UK, NG7 2RD; Department of Oncology, Nottingham University Hospitals NHS Trust, Nottingham, UK, NG7 2UH; School of Veterinary Medicine and Sciences, Faculty of Medicine and Health Sciences, University of Nottingham, Nottingham, UK, LE12 5RD

## Abstract

Retroviruses commonly cause neoplasia in many species but have not been described in guinea pigs. While historical evidence exists for the presence of an active retrovirus in domestic guinea pigs (*Cavia porcella),* there has been no description of the sequence of these viruses or their role in disease in guinea pigs. This paper uses genome mining of the published domestic and wild (*Cavia aperea)* guinea pig genomes for retroviral sequences to identify the gamma and beta-retroviral complement of the *Cavia* genome, describing eight groups of viruses with evidence of recombination between virus groups. The most intact group, gamma-like retroviruses related to HERV-T (integration estimates of 855,000 to 3.8 million years ago), has five near full length loci that are likely capable of active infection. RNA-Scope In-situ Hybridisation of archived formalin-fixed, paraffin embedded (FFPE) guinea pig lymphoma sections with a probe for one of these loci demonstrated viral RNA expression in lymphoma tissue, strengthening the case for a role of these viruses in the high incidence of leukaemia and lymphoma in this species.

## Introduction

Guinea pigs are a popular pet and laboratory model species due to their small size, ease of care and breeding. However, they are not as often presented for individual veterinary care as other species and causes of death are rarely fully investigated. This has led to a dearth of knowledge of guinea pig diseases, a situation that is beginning to be rectified with several post-mortem and surgical biopsy-based studies such as the 2,474 cases in Bertram et al 2025, 493 in Dobromylskyj et al 2023 and 204 in Nešpor et al 2023. These large-scale studies demonstrate a high incidence of neoplasia, with up to 20.5% of all autopsied animals in the Bertram et al (2025) study presenting with one or more neoplastic lesions. Common tumour types include thyroid adenoma/adenocarcinoma, lipoma, mammary adenoma/adenocarcinoma (interestingly as common in males as females), uterine leiomyoma/leiomyosarcoma, hair follicle tumours such as trichoepithelioma and trichofolliculoma and leukaemia/lymphoma. The incidence of leukaemia and lymphoma varies depending on the study, focus population and sampling method (3.4%, 7%, 10.46%) but is rated as one of the five most reported types of neoplasia for this species across all published studies. In comparison, rates of lymphoma are reported as very low in other South American rodents commonly kept as pets, namely chinchillas (*Chinchilla lanigera*) and degus (*Octodon degus*) (Hocker et al. 2017; Jekl, 2020; Mans and Donnelly, 2020). The prognosis for lymphoma in guinea pigs is typically poor (Pignon and Mayer, 2020).

Retroviruses are a group of RNA viruses with an unusual life cycle, in that they create a double stranded DNA copy of themselves that is integrated into the host cell’s DNA. If this cell is the progenitor of a germ line cell the virus will become inherited as part of the DNA of its host during reproduction. These inherited viruses are known as endogenous retroviruses (ERVs), their horizontally infectious counterparts are known as exogenous retroviruses (XRVs). The *Retroviridae* family taxonomy is split into two subfamilies, the *Spumaretrovirinae* (not associated with disease) and the *Orthoretrovirinae.* The *Orthoretrovirinae* are further split into six genera based on sequence similarity and complexity of accessory genes, in addition to the four basic retroviral genes of *gag, pro, pol* and *env* (Black et al. 2025). Of those six genera the most recently actively infectious or endogenizing viruses in mammals are those from the *Gammaretrovirus* and *Betaretrovirus* genera (LaMere et al. 2025).

The process of endogenisation is very common, with ERVs reported in all vertebrates studied to date and ERVs making up 8% of the human and 10% of the house mouse (*Mus musculus*) genomes (Denner, 2010). Most ERVs have been integrated into the genome of their host for millions of years and are mutated and inactive as viruses, though some have been co-opted for cellular functions, such as the syncytins that form placental fusion proteins in many species (Denner, 2016).

While most ERVs are thought of as “fossil” records of retroviral infections earlier in the evolutionary history of a species (Katzourakis and Gifford, 2010), there are several groups of currently evolving gamma and betaretroviruses that are more recently integrated and have infectious counterparts. Examples include Jaagsietke retrovirus (JSRV) in sheep (Armezzani et al. 2014), feline leukaemia virus (FeLV) in cats (Erbeck et al. 2021), koala retrovirus (KoRV) in koalas (Quigley et al. 2021), murine leukemia virus (MuLV) and murine mammary tumour virus (MMTV) in mice (Boeke and Stoye, 1997). Classical XRV infections trigger neoplasia by insertional mutagenesis and disruption of cellular oncogenes, often by stimulating overproduction of cellular genes driven by viral transcriptional promoters. Some viral envelope proteins, such as those of JSRV and some strains of MuLV also stimulate cellular growth (and neoplasia) in addition to insertional mutagenesis (Pederson and Sorensen, 2010).

It is by now well established that in species with recent retroviral integrations both endogenous and exogenous viruses can and do trigger high rates of neoplasia in their hosts, either by the inheritance of a predisposition to neoplasia from ERVs integrated in or near oncogenes or the insertion of new retroviral loci (somatic or endogenous) triggering oncogenesis (Young et al., 2012; McEwen et al., 2021). Mice, koalas and cats all have active, recently integrated gammaretroviral insertions, which cause very high lymphoma and leukaemia rates (Young et al. 2012; Gillet 2014; Chiu et al. 2018). These species exhibit high expression levels of viral RNA in neoplastic tissue (McEwen et al 2021, Chiu et al 2018).

There has long been speculation as to whether there is an underlying retroviral cause for lymphoma in guinea pigs, as a transmissible lymphoma causing inoculate was reported in the 1960’s (Opler, 1967, 1968) and reports of retroviral particles visible with electron microscopy from tissues and cell culture date back to the 1970’s (Hsiung and Kaplow, 1970; Nayak and Murray, 1973; Murray and Nayak, 1974; Rhim et al., 1974; Michalides et al., 1975; Davis et al. 1978; Lerner-Tung et al. 1995). These viruses however could not be sub-cultured and, as it became apparent from further studies that these viruses were endogenous rather than infectious (Nayak, 1974; Davis and Nayak, 1977; Dahlberg et al., 1980), interest in the topic stalled.

However, ERVs in guinea pigs and other South American rodents have not been studied in depth. Recent advances in sequencing technology have also enabled detailed analysis of expression of individual loci. Given the ability of the guinea pig viruses to form virions it is likely (in common with MuLV) that they have the potential to re-integrate into cells triggering cancer. Recently integrated ERVs also have a variety of unpredictable interactions with their infectious counterparts, for example, blockade of infection (JSRV) (Mura et al, 2004), recombination with infectious counterparts enhancing pathogenesis (FeLV B) (Erbeck et al. 2021), the integration of cellular oncogenes which become transmitted as infectious viruses (some MuLV strains) (Pederson and Sorensen, 2010), failure to recognize virus as foreign (due to lifelong infection) and continual viral re-insertion triggering high cancer rates (KoRV) (McEwen et al., 2021) and even resurrection of infectious viruses by recombination of multiple defective genomic copies (MuLV) (Young et al., 2012).

Based on these observations, we hypothesised that intact gammaretroviral ERVs are likely to be present in the guinea pig (domestic and wild, *Cavia porcellus* and *Cavia aperea)* genome and absent in the genomes of degus (*Octodon degus*) and chinchillas *(Cavia lanigera),* and that these ERVs will be highly expressed in haematopoetic neoplasia tissue. Therefore, in this study, we have screened the reference genomes of four South American rodent species and characterised ERV loci containing remnants of the three major retroviral genes - *gag*, *pol* and *env*, in order to identify potentially intact insertions with the potential to cause neoplasia and to further understand the evolutionary history of retroviruses in these species. The most intact insertion identified was a 9,944 bp gammaretrovirus-like locus in the domestic guinea pig which appears to be fully intact. Expression of this locus in FFPE tumour tissues from guinea pigs with a histological diagnosis of lymphoma was then confirmed by RNAscope.

## Results

The two species of *Cavia* - *Cavia porcellus* and *Cavia aperea* for which whole genome sequencing data is available through Ensembl (Dyer et al. 2025) were screened for ERVs alongside two additional Caviomorph species - *Octodon degus* and *Chinchilla lanigera –* (Figure 1A, Supplementary Table 1).

**Figure 1:**
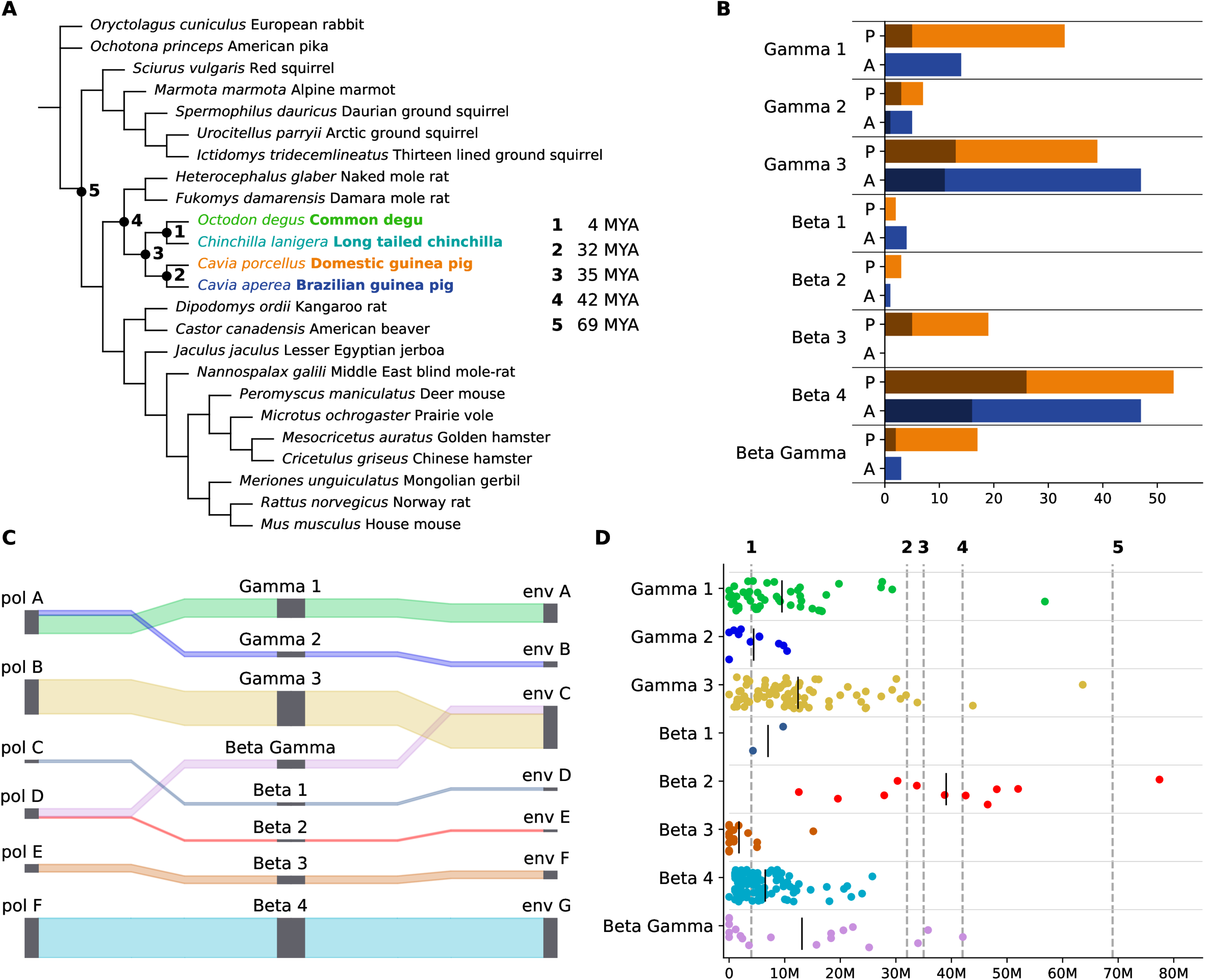
Comparison of Caviomorpha ERVs. A) Rodent phylogenetic tree based on the Ensembl Compara tree (Dyer et al. 2025). Caviomorph species are shown in bold. Node labels are time since the last common ancestor, based on Kumar et al. (2022). B) The number of ERV regions from each cluster identified in each *Cavia* species. Shaded bars show the number of ERV regions with a combined ORF length >6000 nt. P, orange – *Cavia porcellus,* A, blue *– Cavia aperea* C) Sankey plots showing the groupings of the *pol* and *env* genes identified in each cluster. Line heights are proportional to the number of sequences in each group. D) Estimated time since integration, based on LTR divergence, for members of each cluster with recognisable LTRs. Vertical lines represent specific divergence dates in the host phylogeny, labelled as in A).

In total, across the two *Cavia* species, 294 regions were identified with open reading frames (ORFs) ≥200 amino acids with strong similarity to each of the retroviral *gag*, *pol* and *env* genes. Of these, 173 were identified in *C. porcellus* and 121 in *C. aperea* (Figure 1B). All sequences from *Cavia* species were retained, for further analysis for *O. degus* and *C. lanigera* sequences were only retained for analysis if they formed an unambiguous phylogenetic cluster with a *Cavia* ERV gene.

Sequences were considered to be potentially intact if the combined length of the *gag*, *pro, pol* and *env* ORFs was ≥6000 nucleotides, based on the total CDS length of International Committee on Taxonomy of Viruses (ICTV) classified members of the *Gammaretrovirus* and *Betaretrovirus* genera with annotated *gag, pol* and *env* genes, which ranges from 6,609 to 12,642 nt (Black et al. 2025). Eighty two regions in *Cavia* species met these criteria– 54 in *C. porcellus* and 28 in *C. aperea* (Figure 1B). The combined length of *gag*, *pro, pol* and *env* in these regions ranged from 6,009 to 7,179 nt.

Full details of all identified regions are provided in Supplementary Table 2 and further details of each individual domain in Supplementary Table 3.

The identified ERV regions can be divided into three groups of gamma-like ERVs, four groups of beta-like ERVs and a recombinant group with beta- and gamma-like regions (Figure 1C, Figure 2, Figure S1). Recombination appears to be common in this last group, with the *gag* and *pol* genes frequently showing different phylogenetic relationships to the *env* genes. Genome diagrams for all identified ERV regions are provided in the Supplementary Data in the genome_plots directory.

**Figure 2.**
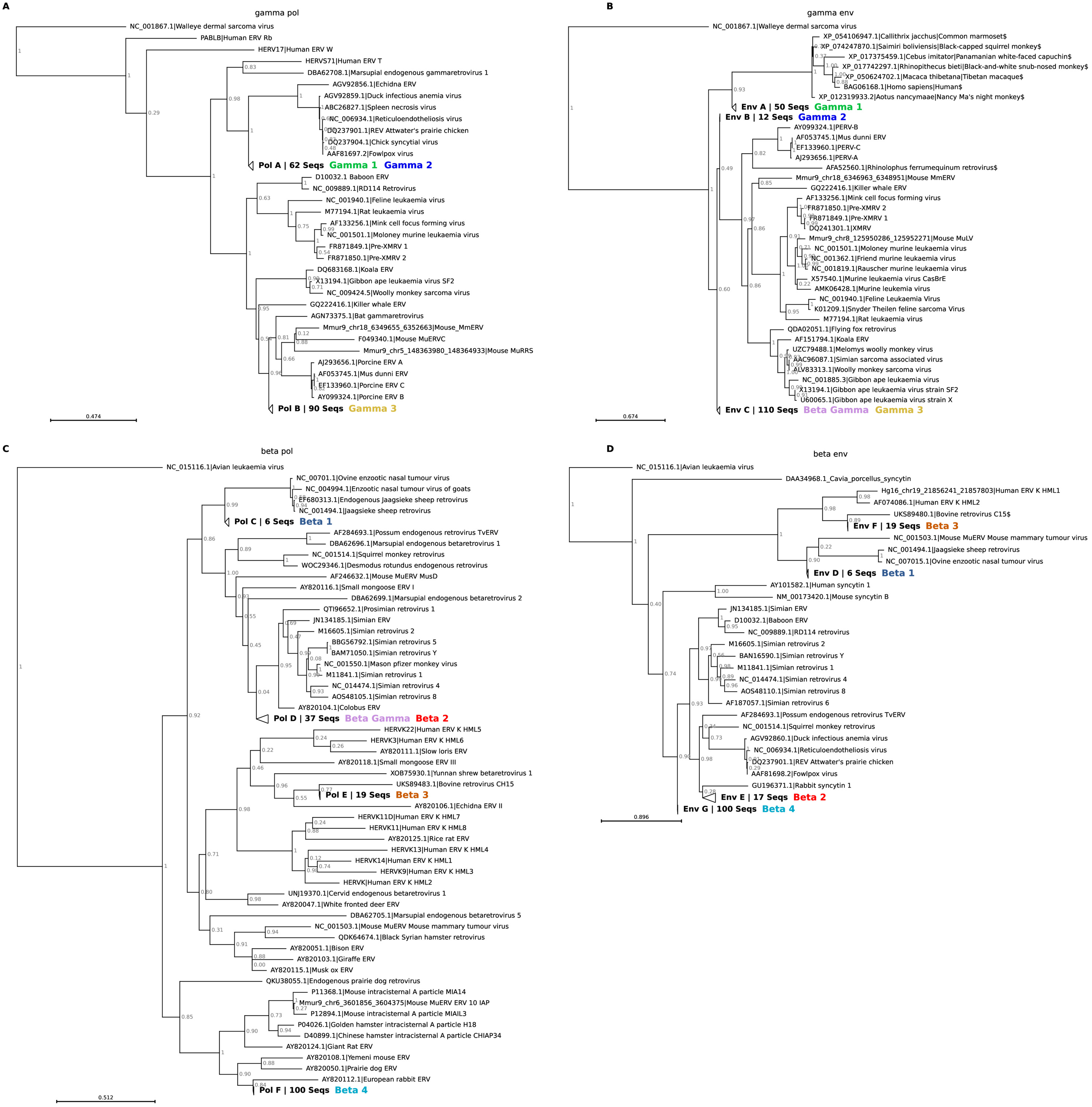
Phylogenetic trees for newly identified Caviomorph ERVs. Maximum likelihood amino acid phylogenetic trees showing the evolutionary relationships of the newly identified ERV ORFs in *Cavia* and other Caviomorphs compared to representative sequences from known exogenous viruses and ERVs for A) gamma-like pol genes, B) gamma-like env genes, C) beta-like pol genes, D) beta-like env genes. Clades containing only a single *pol* or *env* gene group have been collapsed, expanded trees are shown in Figure S1 and full trees are available in the Final_Trees directory in the Supplementary Data. Coloured bold text shows the ERV clusters for which ORFs were identified in each group. Sequences marked with $ are HERV-S71 related sequences from primates identified using BLASTP. Node labels represent branch support.

Six distinct phylogenetic groups of *pol* genes were identified and seven groups of *env* genes, several of which are shared between different clusters (Table 1, Figure 2, Figure S1).

### Gamma-Like Clusters

Clusters were classed as gamma-like if the *pol* and *env* genes both clustered with the *Gammaretrovirus* and gamma-like retroviruses in phylogenetic analysis (Figure 2).

Gamma cluster 1 includes 33 regions in *C. porcellus,* 14 in *C. aperea* and 3 in *O.* degus (Figure 1B). Members of this cluster have *gag-pol* regions which are part of *pol* group A, the same phylogenetic group as members of gamma cluster 2 and the gamma-beta cluster, (Figure 1C, Figure 2). These regions are closely related to, but distinct from, those of reticuloendotheliosis virus (REV, *Gammaretrovirus aviretend*) and its relatives, which include several avian pathogens and endogenous REV-like insertions in the echidna (*Tachyglossus aculeatus*) and the Malagasy carnivore *Galidia elegans* (Niewiadomska and Gifford, 2013), plus human endogenous retrovirus (HERV) T and an endogenous gammaretrovirus identified in Tasmanian devils (*Sarcophilus harrisii*) (Marsupial endogenous gammaretrovirus 1) (Harding et al. 2024). The *env* region forms *env* group A, which is unique to gamma cluster 1. This *env* is related to that of HERV-S71 (Werner et al. 1990) and its relatives (Figure 2B). The most similar *env* region labelled as retroviral in NCBI Protein to the *env* group A consensus is that of Rhinolophus ferrumequinum retrovirus, found in the greater horseshoe bat (Cui et al. 2012), with which it shares only 36% amino acid identity across 389 aa.

Members of gamma cluster 1 are mostly degraded, but five longer insertions were identified, all in *C. porcellus* (Figure 3). All these insertions have recognisable LTRs with >99% sequence identity, and four of the five have LTRs 1,063 to 1,065 nt in length (the other has 825 nt LTRs). Given an estimated mammalian neutral substitution rate of 2.2 × 10⁻⁹ substitutions per site per year (Kumar and Subramanian 2002), the LTR divergence of these five insertions gives estimated integration date of 855,000 to 3.8 million years ago (MYA), after *C. porcellus* and *C. aperea* diverged from their most recent common ancestor an estimated 4 MYA (Figures 1A, D). These insertions, particularly r41 and r157, bear all the hallmarks of active ERVs capable of producing functional viral particles (Figure 3). R41 also has the longest combined ORF length of any of the gammaretroviruses identified in Caviomorphs here, at 7,179 nt. Therefore, these insertions, particularly r41, are strong candidates for producing the viral particles previously observed in guinea pig tissues (Opler, 1967, 1968, Hsiung and Kaplow, 1970; Nayak and Murray, 1973; Murray and Nayak, 1974; Rhim et al., 1974; Michalides et al., 1975; Davis et al. 1978; Lerner-Tung et al. 1995, Nayak, 1974; Davis and Nayak, 1977; Dahlberg et al., 1980)

**Figure 3:**
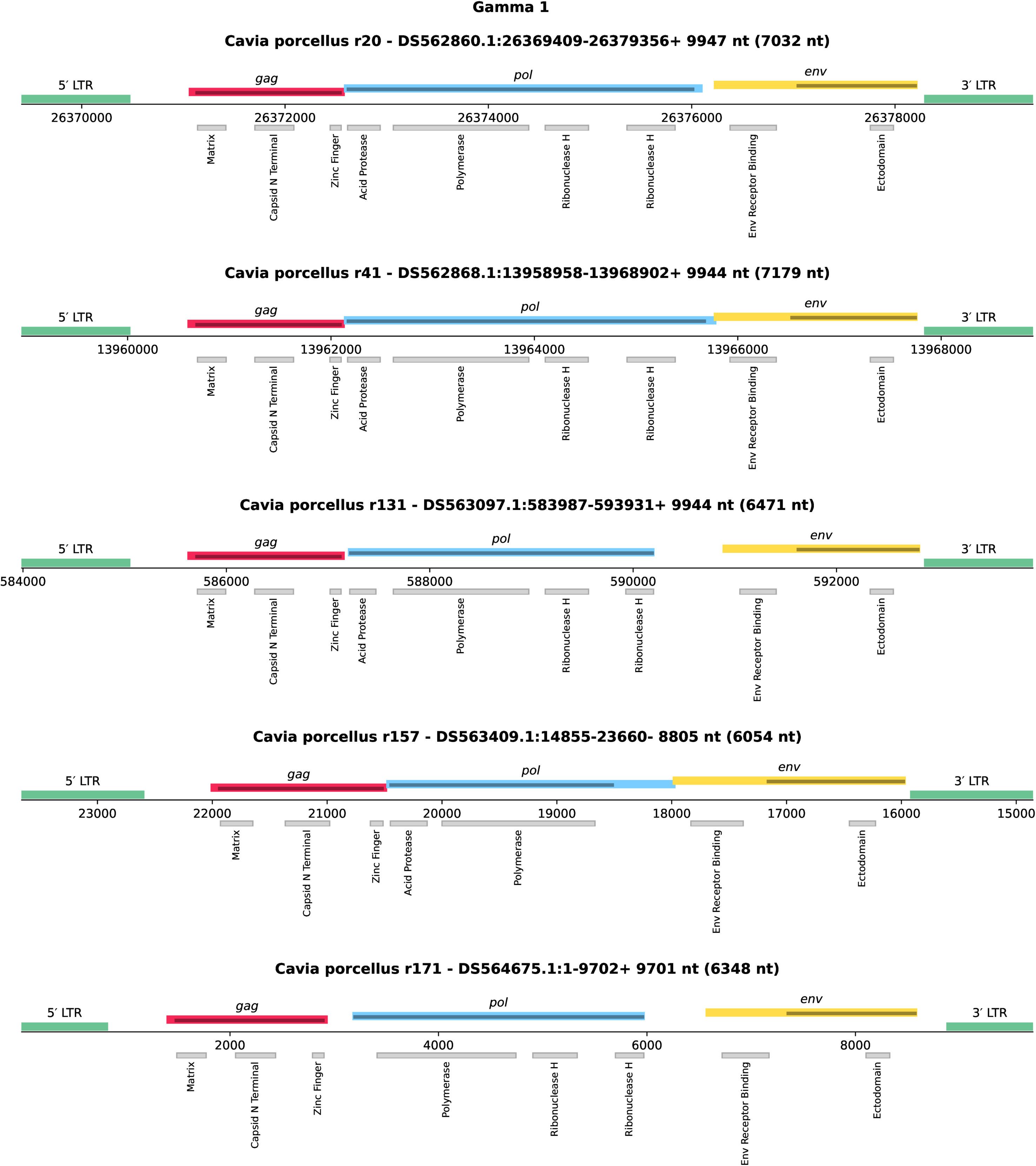
Genome structure of the potentially intact members of gamma cluster 1. Positions are relative to the chromosome or contig in which the region is found. For coding regions, bars above the x-axis show the extent of the ORFs encoding *gag, pol* and *env*, with shaded regions within these bars showing the extent of the domain identified with HMMER. For LTRs, bars above the x-axis show the extent of the regions which could be aligned with BLAST. Bars below the x-axis show protein SUPERFAMILY domains identified within the coding regions, full details are provided in Supplementary Table 3.

Gamma cluster 2 consists of insertions found only in *Cavia –* 7 in *C. porcellus* and 5 in *C. aperea* (Figure 1B). Members of this group again have *pol* genes in *pol* group A, indistinguishable from those of gamma cluster 1, but the *env* genes are unique, forming *env* group B (Figure 1C, Figure 2). The closest relative to the *env* consensus labelled as retroviral in NCBI Protein is from murine leukemia virus (MuLV, *Gammaretrovirus murleu*), but the env gene identity is only 29% across 123 amino acids, making this is the most divergent gamma-like *env* gene identified here. This group is highly intact – four of the 12 insertions have combined ORF lengths >6000nt (Figure 1B). It also appears to have integrated relatively recently compared to the other gamma-like clusters, two of the highly intact insertions in *C. porcellus* (r108 and r114) have no differences between their LTRs, which is a strong indicator of a recent insertion event, and the earliest estimated integration date for a member of this group is 10 MYA, with a mean of 4 MYA (Figure 1D). Members of this group therefore appear to have the potential to produce viral particles.

The third and final group of gamma-like sequences, gamma cluster 3, also has the apparent potential to produce virions. This cluster is widespread in *Cavia* genomes (39 in *C. porcellus* and 47 in *C. aperea*) (Figure 1B). Four insertions were also identified in *O. degus*. Members of this group have a unique group B *pol* gene and a group C *env* gene in common with the beta gamma cluster. Both *pol* and *env* resemble MuLV and related gammaretroviruses such as porcine endogenous retroviruses A to C (PERV-A to PERV-C) and *Mus dunni* ERV (Figure 2A, Table 1). Many members of this group in *Cavia* appear to be potentially intact (Figure 1D). One region (*C. porcellus* r54) has identical LTRs and the mean estimated integration date based on LTR divergence is 12 MYA.

### Beta-like Clusters

Beta cluster 1 members have *pol* genes in the *pol* D cluster and *env* genes in the *env* D cluster (Figure 1C, Figure 2B, D). There are six members of this group, all in *Cavia,* 2 in *C. porcellus* and four in *C. aperea* (Figure 1B). None appear to be intact. Both the *pol* and *env* genes of this cluster are related to a group of pathogens of ruminants - enzootic nasal tumour virus of goats, Jaagsieke sheep retrovirus (*Betaretrovirus ovijaa*) and Ovine enzootic nasal tumour virus.

Beta cluster 2 contains 17 *pol* genes: three in *C. porcellus*, one in *Cavia aperea,* 12 in *O. degus,* and one in *C. lanigera* (Figure 1B). This is the only ERV cluster which had its highest number of repeats in the *O. degus* genome. The *pol* gene is in *pol* group D and is shared with the beta gamma cluster. It resembles the *pol* gene of the simian retroviruses, with maximum similarity to simian retrovirus 1 (64% identity across 535 amino acids) (Figure 1C, Figure 2). The *env* gene is in *env* group E, the closest relative of which is REV and its relatives and an endogenous betaretrovirus initially identified in the possum (*Trichosurus vulpecula*) (Bailie et al. 2010) and recently shown to be widespread in marsupials (Lillie et al. 2024, Harding et al. 2024). None of the ERVs in the cluster met our criteria for being potentially intact (Figure 1D, EB). Based on LTR divergence, this is by far the oldest group of Caviomorph ERVs, with one ERV dated at 77 MYA. This date is older than the most recent common ancestor of the rodents, so it is likely to be an overestimate given the absence of this group in other rodents. However, the presence of members of this group in all four Caviomorph species would be consistent with at least some integration events occurring before the divergence of the *Octodon-Chinchilla* clade from the *Cavia* clade, estimated at 42 MYA (Figure 1A).

Beta cluster 3 has both *pol* and *env* genes clustering with bovine retrovirus CH15, a pathogenic exogenous retrovirus causing neurological disease in cows (Hierweger et al., 2021). Yunnan shrew retrovirus (Feng et al. 2024) also has a related *pol* gene. Members of this group were found exclusively in *C. porcellus,* where 19 copies were identified (Figure 1B). Five of these are potentially intact and appear to have integrated within the last 4 MY, after the divergence of *C. porcellus* and *C. aperea* from their last common ancestor (Figure 1A, D).

Beta cluster 4 is the largest group of ERVs identified here, with 100 representatives, all in *Cavia* species. The *pol* genes of this group (*pol* group F) cluster with the rodent intracisternal A particle genes and with a cluster of ERVs identified in rodents and rabbits (Gifford et al. 2005) (Figure 2B). The *env* genes (*en*v group G) cluster with the simian retroviruses (Figure 2D). Intracisternal A particles do not have *env* genes (Reuss, 1992) but related elements with *env* regions are known, for example in mice (Ribet et al. 2008). Members of beta cluster 4 are highly intact, almost half have combined ORF lengths >6000 nt. Based on LTR divergence, all integrated with the last 32 MY, which is consistent with their absence in *Octodon* and *Chinchilla* species (Figure 1A, D).

### Cross-Genus Recombinants

A few ERVs were identified in *Cavia* with *gag* and *pol* genes from one genus and *env* genes from another, a common phenomenon in retroviruses.

The beta gamma cluster has beta-like *pol* genes and gamma-like *env* genes and consists of 17 regions in *C. porcellus* and 3 regions in *C. aperea* (Figure 1B). The *pol* gene is in *pol* group D and resembles that of the simian retroviruses and beta cluster 2 (Figure 1C, Figure 2C). The *env* gene is in *env* group C and resembles that of MuLV and gamma cluster 3 (Figure 1C, Figure 2B). This group appears to be relatively ancient, with only two potentially intact members and a maximum estimated insertion date of 42 MYA and a mean of 13 MYA; both would be consistent with integration after the divergence of *Cavia* from the other Caviomorphs but before the divergence of *C. porcellus* and *C. aperea* (Figure 1A, D).

### Expression of Gamma Cluster 1

To examine whether the most intact gammaretroviral loci (members of gamma cluster 1) is expressed in guinea pig lymphoma samples from (Dobromylskyj et al 2023), custom RNAScope probes were designed against the most intact locus (R41 locus in Supplementary Table 2). These were applied to five archival FFPE biopsy samples submitted from pet guinea pigs to a UK commercial diagnostic pathology laboratory (case details in Supplementary Table 5). These guinea pigs had all been diagnosed with a solid mass lymphoma (head and neck region), by a board-certified veterinary pathologist. Animals ranged in age from 2 years 7 months to 3 years 9 months, with two samples from guinea pigs of unspecified age. Two of the animals were female, one male and two of unspecified gender.

All five samples were positive with the experimental probe (Figure 4, Figure 5). Negative control probe slides (*Bacillus subtilus* dihydrodipicolinate reductase DapB) were all negative. Four out of five positive control probe slides (*Cavia porcellus* peptidylprolyl isomerase B Ppib) were positive.

**Figure 4:**
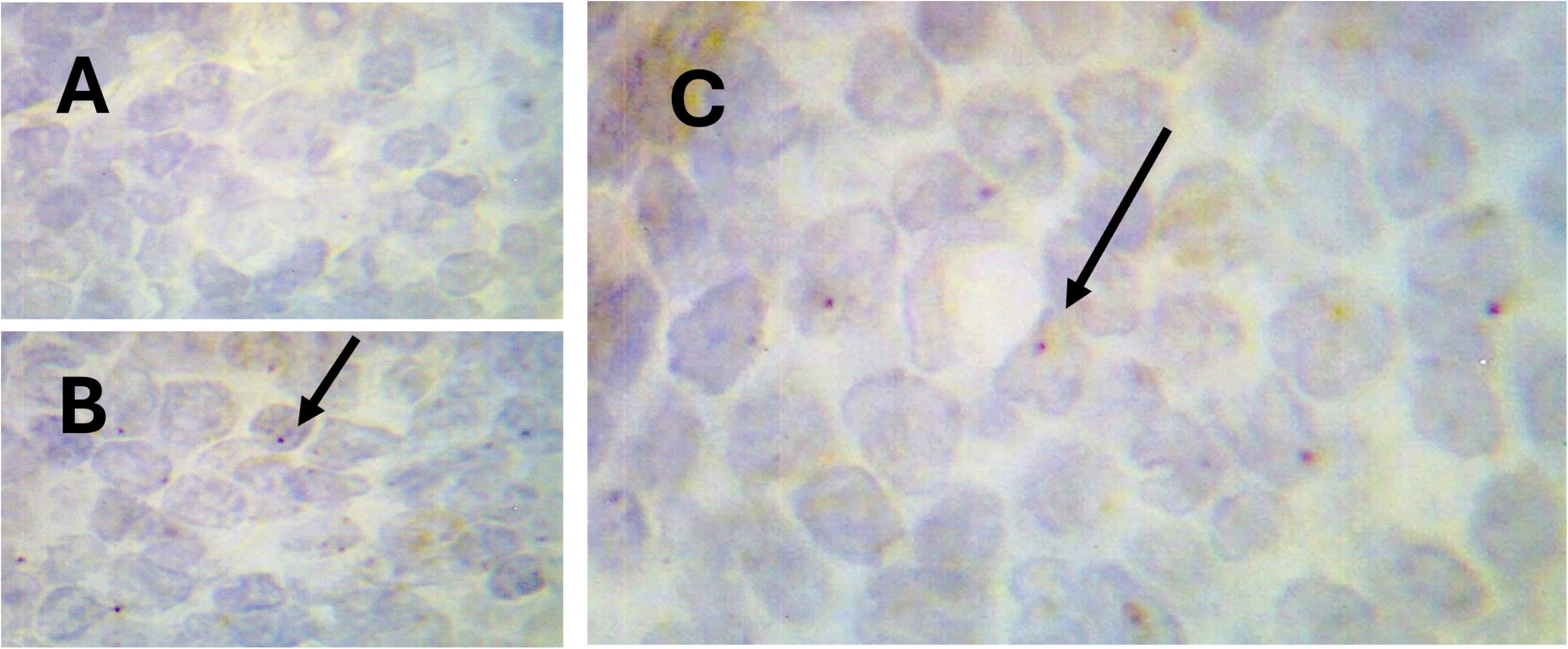
Guinea pig lymphoma sections, case 2. 100x magnification, haematoxylin counterstain, DAB (brown) probe. Arrows = positive staining cells. A= Negative control (*Bacillus subtilus* dihydrodipicolinate reductase DapB), B= positive control (*Cavia porcellus* peptidylprolyl isomerase B Ppib) and C=experimental probe (*Cavia porcellus* Gamma Cluster 1 locus)

**Figure 5:**
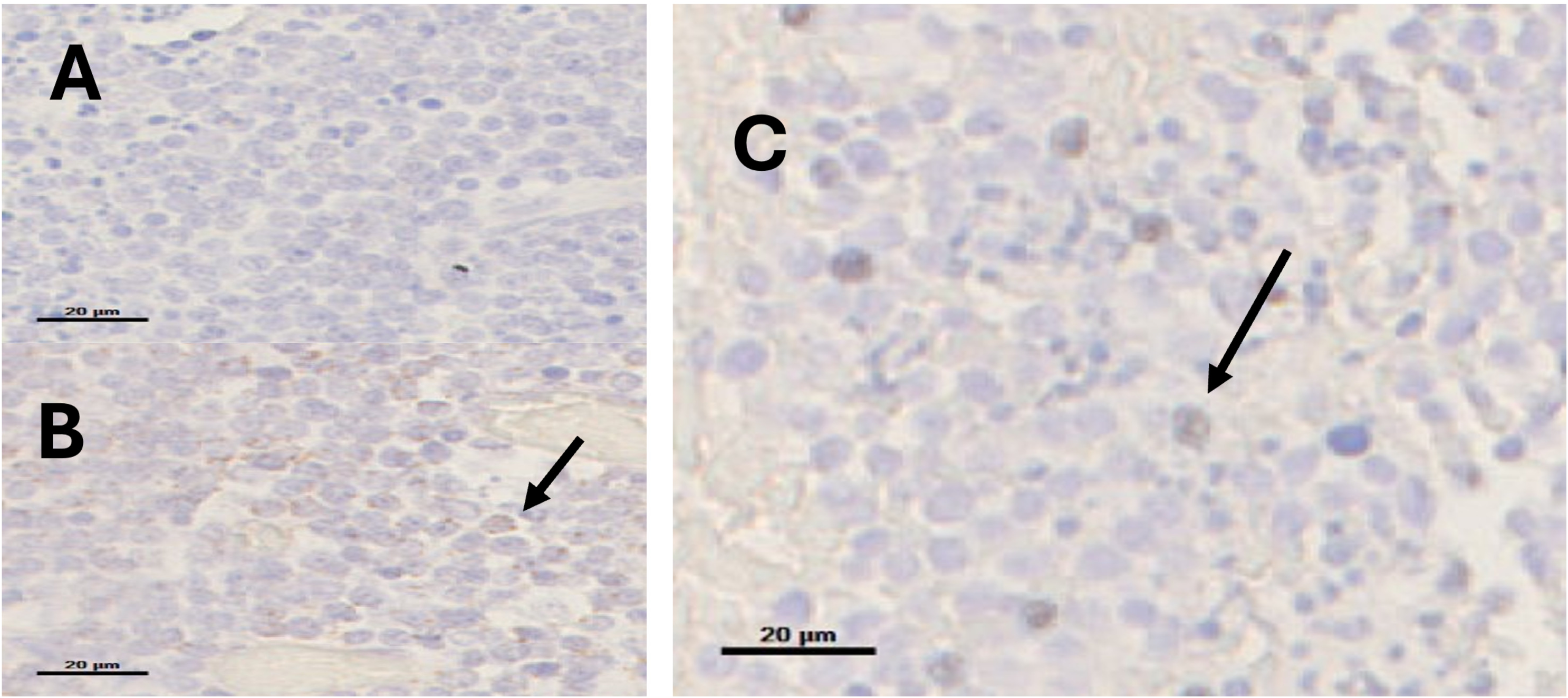
Guinea pig lymphoma sections, case 1. 40x magnification, haematoxylin counterstain, DAB (brown) probe. Arrows = positive staining cells. Scale bar 20 µm A= Negative control (*Bacillus subtilus* dihydrodipicolinate reductase DapB), B= positive control (*Cavia porcellus* peptidylprolyl isomerase B Ppib) and C= experimental probe (*Cavia porcellus* Gamma Cluster 1 locus)

## Discussion

The work presented here provides evidence for a guinea pig specific, recently endogenised gammaretrovirus within the guinea pig genome that has potentially functional loci expressing RNA in lymphoma biopsies from pet animals.

It is not known whether there is an infectious (exogenous) counterpart of this virus currently circulating in guinea pigs or whether the identified gamma cluster 1 loci are polymorphic in the presence/absence of particular loci between individual animals or breeds of guinea pig. However given the high rate of lymphoma in this species (Bertram et al 2025, Dobromylskyj et al 2023 and Nešpor et al 2023) and the overlap in virus and pathology with other species with recently endogenised gammaretroviruses (such as mice, cats and koalas) (Erbeck et al. 2021, Quigley et al. 2021 and Boeke and Stoye, 1997) further investigation of the pathophysiology of these retroviruses in the aetiology of lymphoma in guinea pigs is warranted. Insertional mutagenesis of retroviral loci triggering lymphoma is a well described mechanism of neoplasia in many species. This may be due to new infectious virus insertions, mobilisation of existing ERV loci in somatic cells or an underlying predisposition to certain cancer types due to inheritance of a particular ERV loci in the germ line (or all three) (Young et al. 2012; Gillet 2014; Chiu et al. 2018).

Further work should include qPCR studies and high depth resequencing of both normal and neoplastic guinea pig tissue at RNA and DNA levels as has been done with other species to establish virus load, sequence variation and locus insertion sites (germ line and somatic) (McEwen et al 2021) as well as attempts to isolate infectious virus. All of these methods are more effective with fresh frozen tissue blocks and archive stores of such samples are surprisingly rare for such a common species so this work would likely require planned prospective sample collection for research. Similarly, there are surprisingly few suitable existing data sets in the open access sequence repositories. Most pathology archives (such as the one used in this study) contain historical formalin-fixed, paraffin embedded (FFPE) samples. In the absence of easily available fresh frozen tissue sources we therefore carried out this study using RNAScope to demonstrate RNA expression as proof of principle for future work.

The age and condition of the samples available make it likely that significant RNA degredation has occurred leading to relatively weak signal in our experiments. However, despite this limitation they do unequivocally demonstrate expression of gamma cluster 1 like sequences in lymphoma tissue with confirmation of specificity confirmed by no signal identified in negative control samples. Future investigation of expression in normal and abnormal tissues other than lymphoma and expression at a protein level would also be warranted. This will require the development and characterisation of staining specificity of antibodies to the virus, a task requiring both time and resources. Electron microscopy of guinea pig tissues or cell lines as performed in the original 1970’s studies on this topic would also be helpful to gain further insight but requires fresh tissue specifically stored and processed for this method and cannot be done with the existing pathology archive.

Note that RNAscope probes have a stated specificity of being able to distinguish transcripts with up to 85% homology. The gamma cluster 1 loci have a pairwise homology of between 66.0 – 91.2%, depending on gene segment and locus meaning that some but not all loci would be detected by this probe. This is a common problem with detection of ERV expression where sequence homology means that most detection methods (including qPCR and single end RNAseq) are not able to distinguish with any accuracy which locus of any related group of ERVs is being expressed. In recent years sequencing technology, either Oxford Nanopore long reads or unpaired reads from Illumina paired-end libraries, has enabled this to be done with a higher degree of accuracy, and this is certainly an area of future work (McEwan et al 2021, Tarlinton et al 2022).

Establishing any role of these virus loci in neoplasia in guinea pigs is needed before proceeding to develop an intervention strategy. In terms of animal welfare, the related feline leukaemia virus is controlled with a vaccine, and this would likely be an option for this species. It is also imperative to know whether a mobile genetic element (either ERV or XRV) is present in this common laboratory model species, as this will likely be a serious confounder of guinea-pig models of infectious diseases and neoplasia.

## Materials and Methods

### Genomic screening for potentially functional ERVS

FASTA files for *Cavia porcellus, Cavia aperea, Octodon degus* and *Chinchilla lanigera*, available as part of Ensembl Release 113 (Dyer et al. 2025), were downloaded via FTP. A full list of genome builds used is available in Supplementary Table 1. These files were screened for ERVs using DIAMOND TBLASTX (v2.1.9, Buchfink et al., 2015) with the default settings, using the retrovirus and ERV amino acid sequence database provided with the tool ERVsearch (v1.0.12, Brown, 2020) as a target database. The results were initially filtered to include only results with a BLAST bit score ≥ 200. Hits separated by less than 10,000 nt were then merged into ERV “regions” using BedTools merge (v2.31.1, Quinlan et al., 2010). These regions were also expanded, where possible, to include an additional 2,000 nt on either side. BED and FASTA files of these merged and expanded regions are provided for each host species in the “ERV_regions” directory in the Supplementary Data. All open reading frames ≥200 amino acids were identified in the ERV region FASTA files using orfipy (v0.0.4, Singh and Wurtele, 2021) with ORFs between stops included and allowing ORFs truncated at the 5′ or 3′ end, these are provided in the “original_ORFs” directory in the Supplementary Data. Where the ORFs contained more than three consecutive “X” residues, they were split and terminal “X”s removed, resulting sequences ≥200 amino acids were retained.

To identify ERV domains within these regions, profile HMMs (pHMMs) were built representing retroviral proteins. To create the pHMMs, amino acid sequences from the ERVsearch database described above were separated by genus and gene and aligned using MAFFT with the linsi algorithm (v7.520, Katoh and Standley, 2012). Alignments were cleaned using the CIAlign (Tumescheit et al., 2022) remove divergent (remove_divergent_minperc 0.4), crop divergent (min_prop_nongap 0.1) and remove insertions (insertion_min_size 1, insertion_max_size 5000) functions, the default settings were otherwise used. The cleaned alignments were converted into pHMMs using hmmbuild from HMMER (v3.3.2, Eddy, 2011) and merged into a single file. pHMMs and cleaned alignments are provided in the “HMMs” directory in the Supplementary Data.

The ERV ORF sequences were compared to the pHMMs using hmmscan from HMMER, with the default settings. The results of this search were filtered to keep only sequences with a HMMER domain score ≥ 50 and with a domain envelope (DE) (envelope here referring to HMMER domain envelope rather than retroviral envelope gene) length > 200 amino acids. Only the highest scoring HMMER hit for each gene against each ORF was retained.

Three gene regions were defined as cases where, within an ERV region as defined above, with <10,000 nts separating ERV-like DIAMOND TBLASTX hits, HMMER hits meeting these criteria were identified for each of the retroviral *gag*, *pol* and *env* genes. These were filtered to exclude regions where any two genes were within the same ORF (as where *gag* and *pol* are expressed as a single ORF a ribosomal frameshift occurs between the two genes, so two ORFs would be computationally detected). Regions were also filtered to only include cases where the start position of the genes is ordered as *gag, pol, env* from 5′ to 3′. For phylogenetic analysis, ORF sequences from these regions were trimmed to include only the HMMER region for the relevant gene. BED and translated FASTA files for the resulting sequences are available in the “domains” directory of the Supplementary Data. For reference, each of these domain regions was also compared to the ERVsearch database using BLASTP (v2.14.1+, Camacho et al. 2009, Altschul et al. 1990) these results are included in Supplementary Table 3.

Preliminary phylogenetic analysis was performed by building alignments with the MAFFT linsi algorithm and phylogenetic trees using FastTree2 (v2.1.11, Price et al. 2010), with the default settings. Non-*Cavia* sequences for which no gene clustered unambiguously with a *Cavia* sequence were then removed. Some reference sequences not clustering with the sequences of interest were also removed for clarity. Alignments were rebuilt with the MAFFT with the linsi algorithm and CIAlign was used to trim the edges of the alignment (--crop_divergent --crop_divergent_min_prop_nongap 0.3). Final trees were created using FastTree2 with the default settings. Full phylogenies are available in NEWICK format in the Supplementary Data in the “phylogenies” directory. All trees were visualised with plot_phylo (v0.0.4, github.com/KatyBrown/plot_phylo).

For genomic regions containing correctly orientated ORFs for *gag*, *pol* and *env* meeting the filtering criteria above, genomic co-ordinates for the regions, ORFs and domains, HMMER and BLASTP results and assigned groups are listed in Supplementary Tables 2 and 3.

Consensus sequences for gene clusters were generated using CIAlign with the majority nongap consensus type and insertions >1 nt removed, plus the edges of the alignment trimmed (--remove_insertions --insertion_min_size 1 --crop_divergent --crop_divergent_min_prop_nongap 0.5 --make_consensus --consensus_type majority_nongap). These were compared to sequences labelled as *Retroviridae* (taxonomy ID 11632) in the NCBI non-redundant protein (nr) database using the online server (https://blast.ncbi.nlm.nih.gov/Blast) on 2025-11-20. The full output is included in the Supplementary Data.

*Pro* genes were identified in beta-like insertions (for gamma-like insertions *pro* and *pol* are in the same ORF) by extracting the region between the 3′ end of the *gag* ORF and the 5′ end of the *pol* ORF using BedTools and identifying all ORFs within this region long than 150 amino acids, using orfipy setting as above. These ORFs were then compared to Pfam (Mistry et al. 2021), profile PF00077, Retroviral aspartyl protease, using HMMER as above, and the longest ORF with a HMMER score ≥ 40 retained. HMMER domains were defined as above. These regions are included in Supplementary Table 2.

To identify LTRs, 5000 nt sequences on either side of the ERV region were extracted using BedTools, where possible. Each pair of sequences was compared using BLASTN (v2.14.1+, Camacho et al. 2009, Altschul et al. 1990) and for each pair, the best scoring hit with an alignment length ≥ 200 and a percentage identity ≥ 80, where available, was retained. LTR regions were defined by the co-ordinates of these BLAST hits. Divergence dates were calculated based on an estimated mammalian neutral substitution rate of 2.2 × 10⁻⁹ substitutions per site per year (Kumar and Subramanian 2002). The positions of these regions and estimated dates are included in Supplementary Table 2.

SUPERFAMILY structural domains were identified in all ORFs using the Interproscan online server on 2025-10-28, with the search limited to SUPERFAMILY only. Full details of all results are provided in Supplementary Table 4.

### RNAscope

RNAscope In Situ Hybridisation probes (ZZ probe design for amplification of detection signal for complementary RNA) were designed against the potentially intact gamma cluster 1 locus at the UCSC Cavpor3.0 genome browser scaffold 9 (this position is the reverse complement of the scaffold 13 loci r41) by Biotechne (catalogue number 1247871-C1, Cp retrovirus). Note that RNAscope probes will distinguish transcripts with up to 85% homology and therefore it is likely that multiple gamma cluster 1 loci may be detected with this probe set. The full sequence of the locus targeted is available in (supplementary file 1)

This probe was used alongside a positive control probe Cp-Ppib (Cavia Porcellus peptidylprolyl isomerase B / cyclophilin B catalogue number 471531) and a negative control probe Dap-B (Bacillus subtilus strain SMY methylgloxal synthase (msgA) dihydrodipicolinate reductase (dapB) and YpjD). These are the standard negative control and species-specific positive control for guinea pig RNA detection with RNAscope. These probes were all brown (chromogenic, Diaminobenzidine (DAB) format.

Formalin fixed, paraffin embedded wax blocks of histologically diagnosed guinea pig lymphomas were obtained from archive store from Finn Pathology (UK), these samples are described in the case series described in Dobromylskyj et al 2023. Five head and neck lymphoma tumours collected between 2016 and 2022 were selected (case details in supplementary table 5). Sections were cut fresh before processing and staining as per manufacturer instructions using RNAscope 2.5 High Definition (HD)-Brown assay kit (Biotechne catalogue number 322300) as per manufacturers instructions (standard tissue pre-treatment) with Haematoxylin counterstain.

## Supporting information

Column labels for Supplementary Table 2

Column labels for Supplementary Table 3

## Data Availability

All sequence data used is freely publicly available. Processed data is freely available at https://doi.org/10.5281/zenodo.18224095.

