## Supplementary figures and images for "Expression of Retroviruses in Guinea Pig Lymphomas"

### Supplementary Figure 1

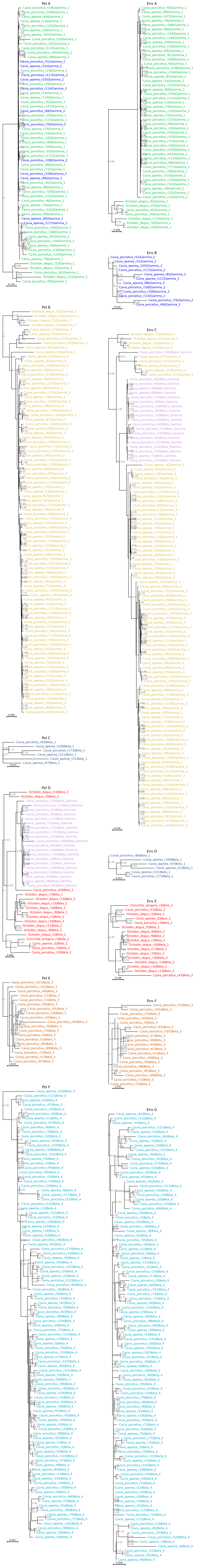
