## Supplementary File Legends for "Expression of Retroviruses in Guinea Pig Lymphomas"

**Figure S1**

Maximum likelihood amino acid phylogenetic trees of the collapsed clades from the phylogenetic trees fromin Figure 2, showing the evolutionary relationships of the newly identified ERV ORFs in Cavia for each pol and env gene type. Full trees are available in the Final_Trees directory in the Supplementary Data. Node labels represent branch support.

**Supplementary Table 1**

Table showing the species names, common names and genome builds for all genomes analysed in this study.

**Supplementary Table 2**

Table showing the details of all ERV regions identified in this study. Columns labels are listed in ST1_columns.tsv.

**Supplementary Table 3**

Table showing details of all individual ERV domains identified in this study. Column labels are listed in ST3_columns.tsv

**Supplementary Table 4**

Table showing the SUPERFAMILY domains identified in the ERV ORFs identified in this study.

**Supplementary Table 5**

Case details of Guinea pig lymphoma sections probed with RNAscope
